# Hollow homes: Roost selection and role of roost morphometrics in the cavity-roosting Bamboo Bat (Genus *Tylonycteris*) from the Western Ghats, southern India

**DOI:** 10.64898/2026.10.20.760418

**Authors:** Sreehari Raman, Baheerathan Murugavel, Mundarappilly R Jagannathan, Anjaly Jose, Unnikrishnan Vadoth Abhishek, Peroth Balakrishnan

**Affiliations:** Department of Wildlife Biology, KSCSTE-Kerala Forest Research Institute, Peechi, 680653, Thrissur, Kerala, India; Department of Wildlife Science, College of Forestry, Kerala Agricultural University, KAU PO, 680656, Thrissur, Kerala, India; Department of Zoology, University of Calicut, Calicut University PO, 673635, Malappuram, Kerala, India

**Keywords:** Bamboo ecosystem, cavity roosting bats, ephemerality, Kerala, specialised roost, Vespertilionidae

## Abstract

Bat species occupy diverse day-roosting habitats, ranging from rock-based to plant-based substrates. Depending on the properties of roosting substrates and the ability of bats to adapt to them, roosts can be stable or ephemeral over time. Bamboo bats (genus *Tylonycteris*) occupy bamboo cavities and possess adaptations such as a flattened skull, thickened thumb pads and modified soles of the feet, making them specialised for roosting within often ephemeral bamboo internodes. The Western Ghats host a bamboo bat species that has traditionally been considered a subspecies of Southeast Asia’s *Tylonycteris pachypus.* However, its precise taxonomic status is unresolved, and recent analyses suggest this population could be a potential new species. Nevertheless, their roosting ecology and the role of bamboo properties in occupancy remain unexplored. By comparing properties of used and unused cavities by the bamboo bats in the Western Ghats, we investigated how physical roost characteristics are associated with cavity occupancy and how the roost properties and space available differed between solitary and non-solitary roosts. We found that only the area of the cavity mouth was associated with occupancy, and all tested properties, including roost volume, were similar for solitary and non-solitary roosts. Our findings are similar to patterns reported from other *Tylonycteris* species, but these findings contribute to the comparative understanding of bamboo bat roost selection across the tropics. It also opens up future avenues to investigate the understudied ecological continuum in the Indian tropics that involves bats, bamboo, pest insects, cavity-dwelling taxa and bamboo harvesters in further detail.

## Introduction

Bats are among the most diverse mammals and occupy microhabitats ranging from rock-based to plant-based substrates, making them useful models for studying habitat selection and social organisation (Gaisler 1979). Roosting substrates are important for bats as they provide shelter and sites for mating, rearing young, and hibernation (Kunz 1982; Swystun et al. 2007; Warakai et al. 2013; Breviglieri and Uieda 2014). While the majority of bat species occupy relatively stable roosting structures, ∼ 30 species are known to roost in ephemeral substrates, including altered tree foliage (tent-making bats; Kunz and McCracken 1996; Arévalo et al. 2026), furled banana leaves (Findley and Wilson 1974; Kunz and Lumsden 2003; Vonhof and Fenton 2004), and unstable tree cavities. The relationship between bats and their roosts can range from obligate to opportunistic depending on roost availability and the ability of bats to adapt to different roosting substrates (Kunz and Lumsden 2003).

Cavity roosting in bats spans this continuum, with species known to occupy both ephemeral and relatively permanent cavities (Kunz and Lumsden 2003). Cavity formation in trees can result from a range of processes including fungal infections, insect or termite attacks, excavation by tree-boring taxa or natural calamities such as lightning strikes or fire (Kunz and Lumsden 2003). Depending on tree species, cavity formation processes and substrate age, some of these cavities may not offer permanency over time and become ephemeral roosting substrates. For bats, such ephemeral cavities can therefore become dynamic resources that require repeated assessment and continued searching. Thus, time invested in roost selection can potentially influence foraging patterns, population dynamics and community structure of cavity-dwelling bats.

Bamboo bats (genus *Tylonycteris*; family Vespertilionidae) are small, insectivorous bats distributed in South and Southeast Asia (Zhang et al. 2005; Wilson and Mittermeier 2019). With adaptations such as a flattened skull, thickened thumb pads and modified soles of the feet, they roost inside cavities of bamboo culms, forming a specialised roosting system (Bates and Harrison 1997; Kunz and Lumsden 2003). The genus comprises six species globally (Simmons and Cirranello 2025), with two reported from mainland India (State of India’s Bats Project Team 2025). The Western Ghats, one of the four biodiversity hotspots in India, hosts a *Tylonycteris* population that was initially considered a subspecies of *T. pachypus,* which is distributed in Southeast Asia (Bates and Harrison 1997; Tu et al. 2017). Based on the biogeography of bamboo bats, Wilson and Mittermeier (2019) described the Western Ghats population as a subspecies of *T. fulvida* that is found in the eastern Himalayas and Indo-Burma biodiversity hotspot, and is referred to as *T. f. aurex*. However, the most recent consensus based on multivariate analysis of this Western Ghats population suggests that it may represent a new species (Chelladurai 2025). Based on this multivariate analysis and the recently annotated regional checklist, we consider the species name to be *T. p. aurex* in this study and use that from here onwards (Raman and Hughes 2020; Raman et al. 2021; Chelladurai 2025). This species also shows specialised adaptations for roosting within bamboo culms (Fig. 1a-d), similar to other *Tylonycteris* species, making it the only bat species currently known to depend exclusively on bamboos for roosting in the Western Ghats. This is also considered rare in this region, with low- to mid-elevation bamboo thickets being their primary roosting habitat (Raman et al. 2021, 2022).

**Figure 1:**
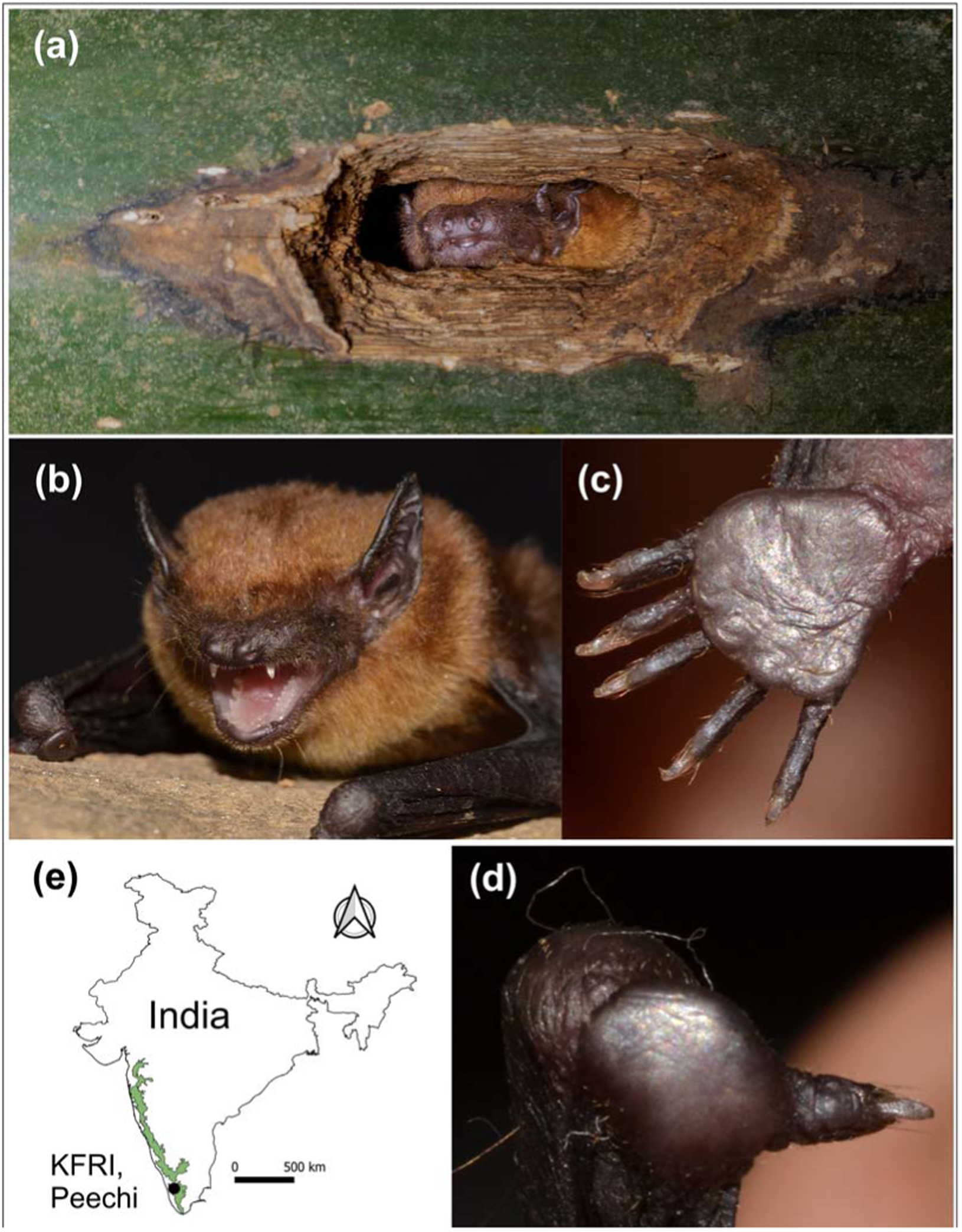
(a) Bamboo bat *Tylonycteris pachypus aurex* roosting inside an insect-made cavity in a bamboo internode; (b) Portrait of a *T. p. aurex* individual showing the flattened head and thick thumb pad; (c) Modified feet with a pad on the sole of the feet. (d) A close-up view of the thumb pad highlighting the adaptation for roosting inside bamboo internodes. (e) Map of India showing our study area with the Western Ghats highlighted in green. Photo credits: (a) Daniel Hargreaves; (b-d) Sreehari Raman

The Western Ghats is home to 28 species of bamboo across seven genera (family Poaceae), five of which are threatened (Bamboo Technical Support Group (BTSG–KFRI)). At least four weevil species (Family Curculionidae) and two leaf beetle species (family Chrysomelidae) are known to attack bamboo during its early growing stages, creating cavities or vertical slits in the internodes (Mathew 2022). Bamboo bats roost inside some of these insect-made cavities and use them for shelter and rearing pups (Medway and Marshall 1972). Being dependent on insect-made cavities in bamboo for roosting and insects for foraging, bamboo bats become a part of a less-explored ecological continuum in this tropical habitat involving bamboo, insects, and bats. However, knowledge on their behavioural ecology, roosting ecology and sociality is limited, particularly in South Asia. This data deficiency on a potentially distinct species not only hinders threat assessment and planning conservation strategies but also creates a knowledge gap in the overall understanding of the bamboo bats, a specialised group of bats in the Paleotropics.

Roost site selection in two sympatric *Tylonycteris* species, *T. pachypus* and *T. robustula,* from Malaysia suggests that the dimensions of the cavity mouth were critical determinants of species occupancy, with the lower limit of the cavity width being constrained by the size of the species (Medway and Marshall 1970). The authors also report the excess availability of suitable roosting sites in their habitat, but they did not systematically measure properties of unoccupied cavities to test whether occupancy was linked with cavity dimensions or simply an outcome of availability. Investigation of roosting associations and group composition in *T. pachypus* and *T. robustula* in Malaysia suggested that the number of bats per roosting group was limited by factors other than available space within the roosts (Medway and Marshall 1972). Selecting a suitable cavity to roost can therefore be a crucial decision for these bats as bamboo cavities are relatively unstable roosting substrates. Cavity properties are dynamic and are likely influenced by the fast growth rates of bamboo shoots, ageing of the culms and external seasonal factors such as monsoons and winds. Not all available cavities might be suitable for roosting, making it important to understand which bamboo or cavity properties are associated with bat occupancy.

The Indian population of *T. p. aurex* does not share its habitat with any other bamboo bat species (Wilson and Mittermeier 2019; Raman et al. 2021), and the factors associated with roost selection in this population remain poorly understood. It is not known whether the roosting associations, group composition and cavity occupancy patterns of other *Tylonycteris* species can be generalised to this population (Medway and Marshall 1970, 1972). Specifically, understanding the associations between bats and bamboo requires a systematic comparison of the properties of occupied and unoccupied bamboo cavities.

We investigated how physical roost characteristics are associated with cavity occupancy in the bamboo bat *T. p. aurex* by comparing properties of used and unused cavities in bamboo culms. Since the hollow bamboo internodes with cavities can be vulnerable to harsh weather and offer space for cavity-dwelling taxa such as insects, arachnids and reptiles, we expected occupied cavities to be associated with properties that offer protection and minimise disturbances from other cavity-dwelling taxa or climatic conditions. We also examined how the physical dimensions and space available differed between solitary and non-solitary roosts. Here we expected no difference between the solitary and non-solitary groups, as reported from the two sympatric *Tylonycteris* species in Malaysia (Medway and Marshall 1972).

## Materials and methods

### Study site

We conducted this study at the Kerala Forest Research Institute (KFRI) campus, Peechi, Kerala. KFRI is situated within a 28-ha reserve forest area, adjoining Peechi-Vazhani Wildlife Sanctuary (Fig. 1e). The area is a part of the Anamalai landscape of southern Western Ghats, lies between 10°31′–10°31′ N latitude and 76°20′–76°21′ E longitude with elevations ranging from 90 to 110 msl. The area comprises tropical moist semi-evergreen and moist deciduous forests, with a diverse assemblage of native trees and bamboos, providing a rich habitat for biodiversity.

### Roost surveys

Thorough daytime surveys were conducted searching for cavities in bamboo thickets located within the KFRI campus from September to December 2025. Cavities were visually inspected for the presence of bats using an LED torch with a red filter. If a cavity hosted bats, the roosting individuals could be seen at or near the cavity entrance during the daytime (Fig. 1a). Such cavities were marked as occupied roosts and were further confirmed by direct observations at each cluster in the subsequent evenings for recording emerging bats. Once located, each of these cavities was given a unique identification code, and information such as bamboo species, cluster size (number of culms in a cluster), and number of culms with cavities (occupied and unoccupied by bats) were recorded. Subsequently, parameters such as distance from ground, angle of inclination, orientation of the cavity mouth, number of cavities in an internode, maximum length of the cavity, maximum width of the cavity, internodal distance, distance from bottom internode to base of the cavity, and distance from top internode to cavity were recorded for all occupied cavities (n=22). To understand roost selection, the same information was also collected from 21 randomly selected unoccupied cavities from the same or adjacent culms. In all bamboo clusters, upward-facing canopy photographs were taken on an Android phone (version 13 TKQ1.221013.002) to estimate canopy openness. During the course of these surveys, some occupied cavities were subsequently abandoned, and one cavity was newly identified. Properties of all these cavities were recorded and incorporated in the analysis as appropriate.

### Roost morphometrics

Cavity metrics such as cavity width and cavity length were measured using digital callipers of 0.01mm accuracy. Distance from the lower and upper nodes (to the nearest margin of the cavity mouth), as well as length and girth of the internode, were measured using a measuring tape. Cavity height above ground was determined using a laser range finder (Bosch GLM 40; Measuring range: 40 m; accuracy: ±2 mm). Orientation of the cavity mouth and angle of inclination with respect to the ground were measured using iPhone applications ‘Compass’ and ‘Measure’, respectively. Culm wall thickness was measured from five randomly selected culms per cluster, and the mean thickness for each cluster was taken as a constant for volume estimation; the volume was calculated using the equation

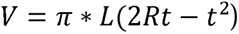

Where ‘L’ is the length of the internode, ‘R’ is the outer radius and ‘t’ is the mean wall thickness of the respective cluster.

For estimating cavity mouth area, we took images of the cavity opening with a ruler and analysed them using ImageJ (Schneider et al. 2012). In every image, scales were set based on calibration from a 4 cm segment of the ruler in ImageJ. The cavity mouth area was delineated using threshold adjustment to isolate the cavity opening, and it was converted to a binary image. Noise and outliers were removed using the brush tool to refine the region of interest (ROI) when required. Selected ROIs were stored in the ROI manager to preserve region boundaries for verification and repeatability. All ROIs were saved for later validation. Cavity mouth area was calculated (measured in cm^2^) via the ImageJ measurement function. Processed images were archived in TIFF format for subsequent reanalysis if required.

Canopy openness was estimated from the photographs using image analysis in R (R Core Team 2025). Images were imported and converted to grayscale using the *EBImage* package (Pau et al. 2010). Sky and vegetation pixels were classified using an intensity threshold selected based on visual inspection of the images. Pixels with a brightness value of >0.7 were considered sky and those with ≤ 0.7 as vegetation. The same threshold was applied consistently across all images. Canopy openness was then calculated as the proportion of sky pixels with respect to the total number of pixels in the image, expressed as a percentage.

### Solitary and non-solitary roost classifications

The occupied cavities were classified as solitary and non-solitary roosts based on the video footage from a separate monitoring study (Murugavel et al., in prep). At each occupied cavity (n=19), a CCTV camera (Model: TrueView 4G) was mounted on a tripod positioned approximately 1.0 - 1.5 m from the cavity entrance and was continuously recorded from 17:30 to 07:30 hrs. From the recordings of entries and exits, it was clear that some cavities were occupied by a single individual during the night and the others had multiple individuals. However, it was not possible to count the exact number of individuals for cavities occupied by multiple bats. Hence, we categorised roosts as solitary or non-solitary and compared their roost properties between these categories.

### Data transformation and statistics

Cavity position in a given internode was estimated as the ratio between the distance of the cavity from the bottom internode to the base of the cavity and the sum of distances from the bottom and top of the internode. This value ranged from 0 to 1, where 0 means the cavity is located at the base of the internode and 1 means the cavity is located exactly on the top of the internode. Orientation of the cavities was recorded in degrees (0-360°). We linearised this circular variable by converting the degrees into radians and used its sine and cosine components in our analysis. These linear predictors captured both north-south (sin) and east-west (cos) orientations, respectively.

All analyses were performed in R (R Core Team 2025). We fitted a generalised linear model (GLM) with a binomial error function and logit link function. Cavity occupancy was used as a binary response variable. Cavity position ratio, orientation (transformed into sin and cosine components), area of the cavity mouth, angle of inclination and height of the cavity from ground level were used as fixed effects. Initial models included Culm ID, Cluster ID and bamboo species as random effects using *lme4* package (Bates et al. 2015) but were subsequently dropped as their variance estimates collapsed to zero (singular fit). Area of the cavity mouth was used as a single measure of cavity size instead of including the maximum cavity length and width as separate predictors. All candidate predictors were checked for multicollinearity using the *performance* package (Lüdecke et al. 2021). Girth of the internode, volume of the cavity, canopy openness, number of culms in a cluster and length of the internode were removed as they had a high (>10) Variance Inflation Factor (VIF).

To investigate roost properties associated with solitary and non-solitary roosts, we fitted another GLM with a binary error distribution and logit link function. We excluded the abandoned roosts and used only the occupied cavities (n=19) for this analysis, with the assigned roost category (solitary and non-solitary) being a binary response variable. Cavity mouth area, roost volume, angle of inclination and height from the ground were used as predictors, as these four properties are likely to influence a roost being solitary or not.

Collinearity among these factors was assessed using VIF and pairwise correlations. All variables had a VIF <2, indicating they captured independent structural aspects and can be included simultaneously in the model. Our global model contained all four predictors, and the full set of candidate models was generated using the ‘dredge’ function in the *MuMIn* package (Bartoń 2025). All candidate models were then ranked based on the Akaike Information Criterion (AICc). As models with ΔAICc <2 are considered to be the best fits, we performed model averaging (‘model.avg’ function in *MuMIn* package) on such models from our set. We therefore report conditional-averaged coefficients and predictor-importance values from summed Akaike weights, allowing inferences about the relative importance of cavity properties for roost preferences. These model results were also cross-verified using univariate non-parametric Mann-Whitney U tests to compare the properties of solitary and non-solitary roosts for all individual predictors, as our dataset had a low sample size.

## Results

### Occupancy

Our initial survey revealed 21 cavities in three bamboo clusters, of which 17 were observed in two independent *Bambusa balcooa* clusters and four in a *Bambusa vulgaris* cluster. The two *B. balcooa* clusters contained 148 and 48 culms, and the *B. vulgaris cluster* contained 123 culms. Bats had occupied cavities in these two bamboo species irrespective of their cluster size (Fig.2a). Within clusters, some culms in *B. balcooa* contained more than one cavity that had bats, whereas no such pattern was seen in the *B. vulgaris* culms (Fig. 2b).

**Figure 2:**
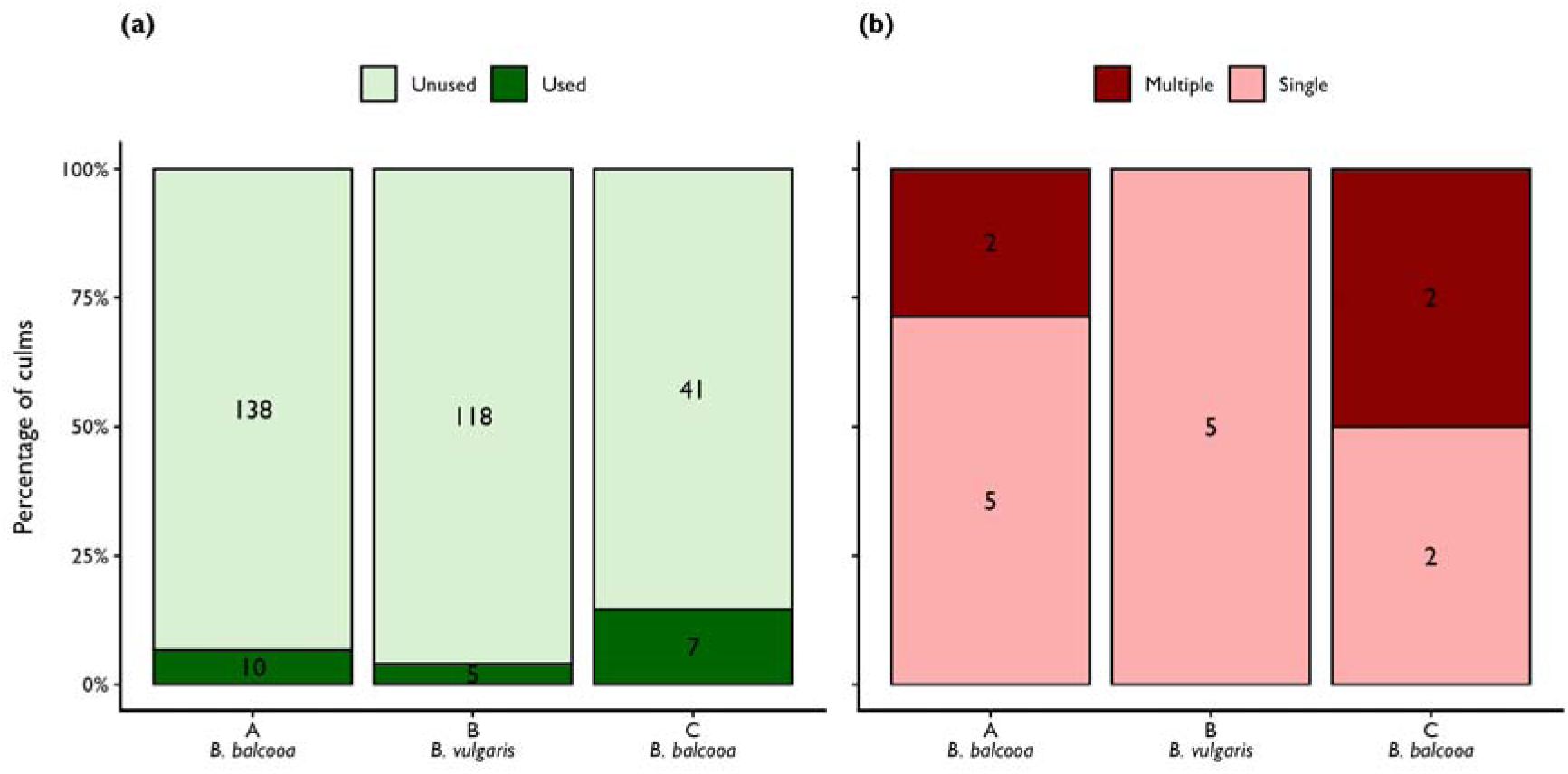
Cavity usage by bamboo bats. (a) among and (b) within culms in the three bamboo clusters in the study area. The number of culms in each cluster is denoted within the bars. The percentage of culms that had cavities with bats was 7%, 4% and 15% (dark green) relative to the overall cluster size in the three clusters. Both the *B. balcooa* clusters had culms with multiple cavities in the same culm occupied by bats (dark red).

During the course of our study, two cavities from *B. balcooa* and one cavity from the *B. vulgaris* cluster were abandoned, and one new cavity was occupied in a *B. vulgaris* cluster (Fig.2a).

### Roost selection

The GLM for roost selection revealed that only cavity mouth area was significantly associated with bat occupancy (β = 1.94 ± 0.97 SE, p = 0.045; Fig. 2), and all the other predictors had no significant associations (Table 1; Fig. 3). On average, the occupied cavities had a larger mouth area (mean ± SD: 1.53 ± 0.57 cm^2^; range: 0.87 to 3.16 cm^2^) than the unoccupied cavities (mean ± SD: 1.08± 0.43 cm^2^; range: 0.58 to 2.54 cm^2^). The positive model estimate and an odds ratio of 6.98 per cm^2^ indicated that cavities with larger mouth area had higher odds of bat occupancy within the sampled range of roosting properties.

**Figure 3:**
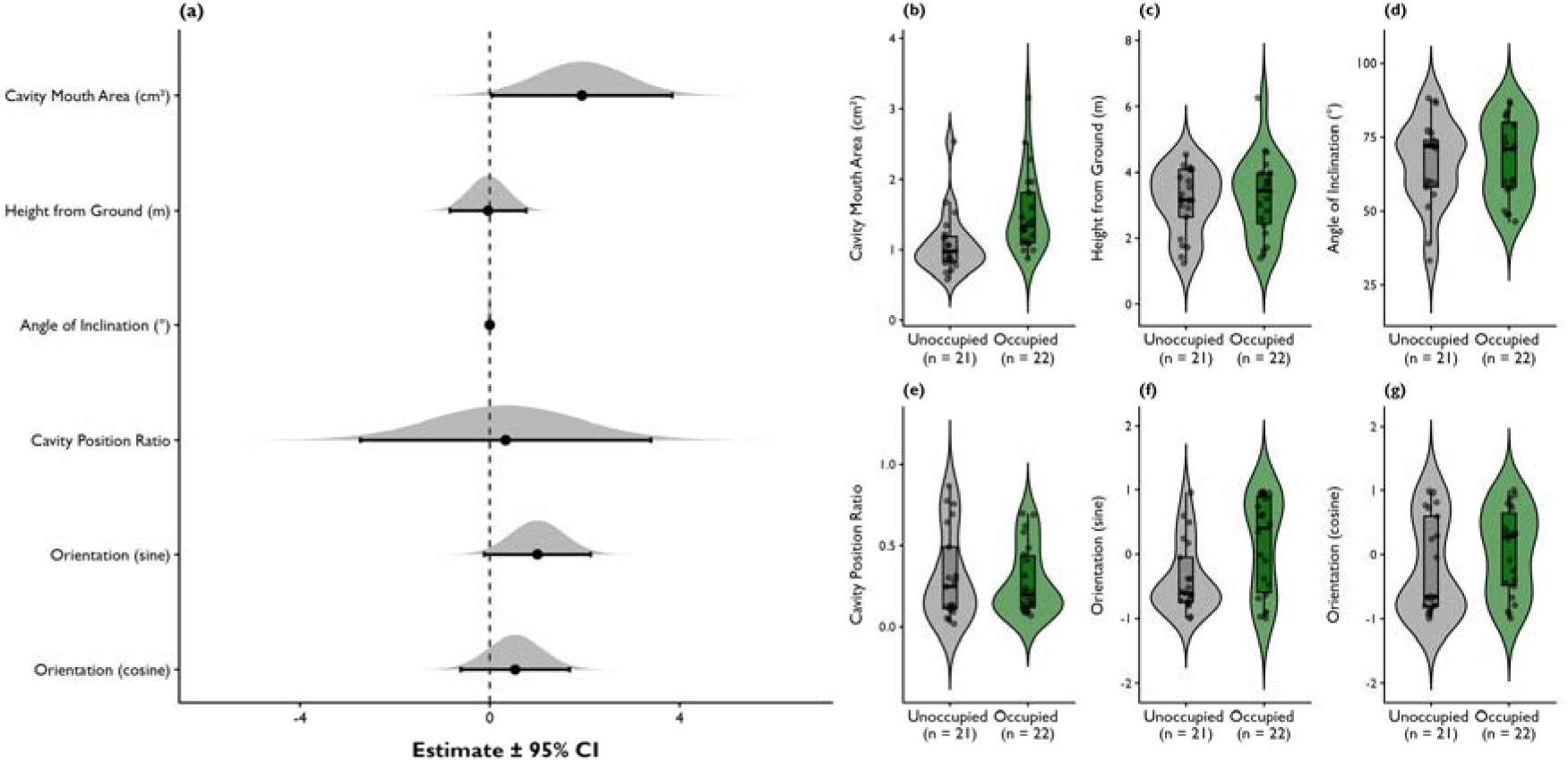
Roost selection and associated properties. (a) The estimated log-odds for the fixed effects from the GLM for cavity occupancy. Points denote the estimated effects, and the error bars show Wald 95% confidence intervals. The dashed vertical line at zero indicates no effect. Positive values indicate higher odds of occupancy, and only the cavity mouth area showed a significant positive association with bat occupancy. (b-g) Distributions of the six roost parameters used as predictors in the GLM comparing unoccupied and occupied bamboo cavities. Each parameter was measured from 22 occupied roosting and 21 unoccupied cavities; sample sizes are denoted as n on the x-axis.

**Table 1:** Odds ratios from the final binomial GLM for roost selection. Roost occupancy was the binary response variable, and the area of the cavity mouth, height of cavities from the ground, angle of inclination, the cavity position ratio, and orientation transformed to sine and cosine functions were candidate predictors. Only the cavity mouth area had a significant association with roost occupancy and is highlighted in bold.

| Predictor | Odds Ratio | CI lower | CI upper | SE | z | p value |
| --- | --- | --- | --- | --- | --- | --- |
| Intercept | 0.103 | <0.001 | 32.111 | 2.929 | -0.775 | 0.438 |
| **Area of the cavity mouth (cm<sup>2</sup>)** | **6.975** | **1.043** | **46.637** | **0.969** | **2.004** | **0.045** |
| Height from the ground (m) | 0.967 | 0.433 | 2.157 | 0.409 | -0.083 | 0.934 |
| Angle of inclination (°) | 1.001 | 0.940 | 1.065 | 0.032 | 0.017 | 0.986 |
| Cavity position ratio | 1.400 | 0.065 | 29.990 | 1.564 | 0.215 | 0.830 |
| Orientation (sine) | 2.739 | 0.893 | 8.403 | 0.572 | 1.762 | 0.078 |
| Orientation (cosine) | 1.710 | 0.543 | 5.384 | 0.585 | 0.917 | 0.359 |

However, the cavity with the largest mouth area in our dataset (3.16 cm^2^) was also one of the three roosts that were subsequently abandoned during the course of our study.

Overdispersion checks based on residual deviance showed no evidence of overdispersion in the model (residual deviance = 46.082, 36 degrees of freedom; dispersion ratio = 1.041, p = 0.824).

### Solitary and non-solitary roosting properties

The global model showed no significant associations between the four tested predictors and an occupied cavity being solitary or non-solitary (Fig. 4; Supplementary Table 1). The model selection approach based on AICc also provided no support for individual predictors, as the null model was the best fit with the lowest AICc (AICc = 24.1, weight = 0.216), followed by the model containing only roost volume (AICc = 24.1, weight = 0.213). However, models including height from ground level and angle of inclination also had ΔAIC <2, suggesting an uncertainty in model selection (Table 2). The conditional model averaged estimates of roost volume (β = 4.29 ± 3.27 adjusted SE, p = 0.188), height from ground level (β = 0.17± 0.66 adjusted SE, p = 0.188), and angle of inclination (β = −0.03 ± 0.04 adjusted SE, p = 0.490) also did not show any clear associations between the roost properties and the roost being solitary or not. This was further supported by the univariate Mann-Whitney U tests, as none of the four predictors showed any significant difference for solitary and non-solitary roosts (Supplementary Table 2). Overall, both multivariate model selection and univariate non-parametric approaches provided little evidence that the measured roost properties were associated with the cavities being solitary or non-solitary roosts.

**Figure 4:**
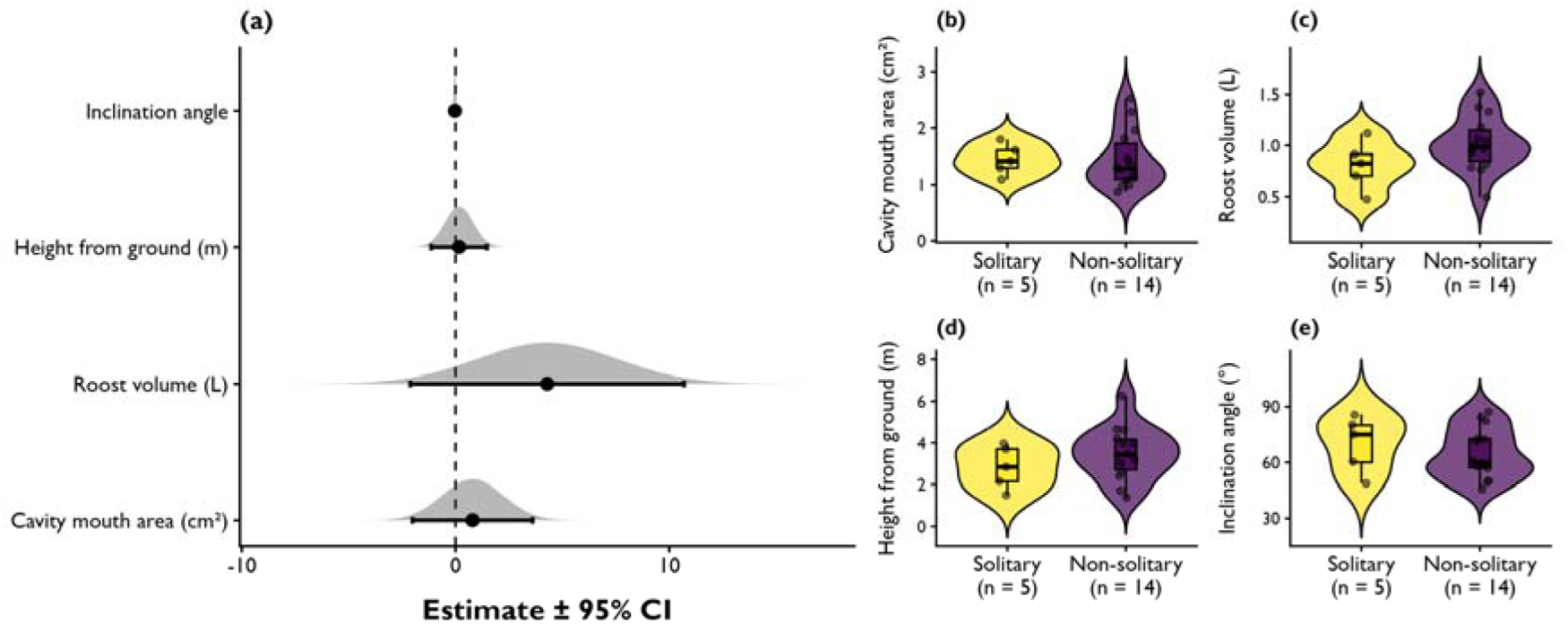
Roost properties associated with solitary and non-solitary roosting. (a) Estimated log-odds for the predictors on roost preference from the GLM. Points denote the estimated effects, and the error bars show Wald 95% confidence intervals. The dashed vertical line at zero indicates no effect. Positive values indicate higher odds of a cavity being non-solitary than solitary, and all the tested predictors showed no significant association with roost preference. (b-e) Distributions of the four roost parameters used as predictors in the GLM comparing solitary and non-solitary roosting cavities. Among the studied roosts, five were solitary, and 14 were non-solitary, and sample sizes are denoted as n on the x-axis.

**Table 2:** Comparison of binomial GLMs for roost preference in bamboo bats using AICc dredge analysis. Roost category (solitary or non-solitary) was the binary response variable. The global model contained inclination angle, cavity mouth area, height of cavity from the ground and roost volume as predictors. Models were selected based on ΔAICc and are ranked from the lowest to the highest AICc. All models with ΔAICc <2 are highlighted in bold.

| Models | df | logLik | AICc | delta | weight |
| --- | --- | --- | --- | --- | --- |
| **Intercept only** | **1** | **-10.95** | **24.136** | **0** | **0.216** |
| **Roost volume (L)** | **2** | **-9.706** | **24.161** | **0.025** | **0.213** |
| **Height from ground (m)** | **2** | **-10.356** | **25.461** | **1.325** | **0.111** |
| **Inclination angle** | **2** | **-10.666** | **26.081** | **1.945** | **0.081** |
| Inclination angle + Roost volume (L) | 3 | -9.446 | 26.491 | 2.355 | 0.066 |
| Height from ground (m) + Roost volume (L) | 3 | -9.451 | 26.502 | 2.366 | 0.066 |
| Cavity mouth area (cm<sup>2</sup>) | 2 | -10.950 | 26.650 | 2.514 | 0.061 |
| Cavity mouth area (cm<sup>2</sup>) + Roost volume (L) | 3 | -9.559 | 26.719 | 2.583 | 0.059 |
| Cavity mouth area (cm<sup>2</sup>) + Height from ground (m) | 3 | -10.344 | 28.288 | 4.152 | 0.027 |
| Inclination angle + Height from ground (m) | 3 | -10.347 | 28.293 | 4.157 | 0.027 |
| Inclination angle + Cavity mouth area (cm<sup>2</sup>) | 3 | -10.665 | 28.931 | 4.795 | 0.020 |
| Inclination angle + Cavity mouth area (cm<sup>2</sup>) + Roost volume (L) | 4 | -9.229 | 29.315 | 5.179 | 0.016 |
| Cavity mouth area (cm<sup>2</sup>) + Height from ground (m) + Roost volume (L) | 4 | -9.354 | 29.565 | 5.429 | 0.014 |
| Inclination angle + Height from ground (m) + Roost volume (L) | 4 | -9.360 | 29.578 | 5.442 | 0.014 |
| Inclination angle + Cavity mouth area (cm<sup>2</sup>) + Height from ground (m) | 4 | -10.337 | 31.531 | 7.395 | 0.005 |
| Inclination angle + Cavity mouth area (cm<sup>2</sup>) + Height from ground (m) + Roost volume (L) | 5 | -9.198 | 33.011 | 8.875 | 0.003 |

## Discussion

By comparing the physical properties of bamboo cavities, we show how roost morphometrics are associated with cavity occupancy of *T. p. aurex* from the Western Ghats. We found that only the area of the cavity mouth was associated with a cavity being occupied by bats. Also, as expected and reported from other *Tylonycteris* species, roost volume or any other tested property was not associated with a roost being occupied by single or multiple individuals (Medway and Marshall 1970, 1972). Our results are based on a systematic comparison of the properties of available unoccupied and occupied bamboo cavities at the same time and provide a better comparative understanding of why bamboo bats choose a cavity.

### Ecology of roost availability

Ecologically, these cavities are made available to the bats by a community of insects that use bamboo shoots at different stages, and these are often recorded as bamboo pests. Bamboos are vulnerable to different pest insects at different growth stages (Shu and Wang 2015). In Malaysia, *Tylonycteris* bats inhabited cavities made by the boring beetle *Lasiochila goryi* (Chrysomelidae: Coleoptera), and the cavity width was defined by the girth of the beetle larva as it bores into the culm wall shortly before pupation (Medway and Marshall 1970). In our landscape, weevils (Curculionidae: Coleoptera) such as *Cyrtotrachelus buqueti*, *Cyrtotrachelus longimanus*, *Cyrtotrachelus dux*, and the hispine borer *Estigmina chinensis* (Chrysomelidae: Coleoptera) are identified as the major pests infesting bamboo shoots and creating cavities (Mathew 2022). Adult female weevils usually drill holes in newly emerging bamboo shoots for oviposition. Once hatched, the larvae (grubs) bore deep into the growing shoots to feed on the inner parenchyma tissue. The initial damage created by primary borers often invites secondary borers, such as *Myocalandra exarata* larvae (Curculionidae: Coleoptera), to further utilise the live bamboo shoots (Mathew 2022). As the surviving shoot matures, these hollowed regions expand into permanent internodal cavities, providing space for cavity-dwelling small vertebrates and invertebrates to shelter and to reproduce (Kovac and Streit 1996; Seshadri et al. 2015; Seshadri and Bickford 2018).

Some of these beetles are known to be seasonal, with the oviposition, larval and pupal stages restricted to the monsoons (Joshi et al. 2008). We could not identify the pest that was involved in the formation of cavities at our study sites, as the study was conducted in the post-monsoon season. However, we observed greater infestation levels in low-density *B. balcooa* clusters compared to high-density *B. balcooa* or *B. vulgaris*. We also found that 17 out of the 22 occupied cavities were in *B. balcooa* (Fig. 2a), and we did not find any culms in *B. vulgaris* with multiple bat roosts (Fig. 2b). Within a radius of 500 – 1000 m from our observed roosting clusters, we find a range of germplasm collection of bamboo species such as*, Bambusa tulda, Dendrocalamus giganteus, Pseudoxytenanthera bourdillonii, P. stocksii, P. ritcheyi, P. madhavii, Thyrsostachys oliveri* and *Schizostachyum brachycladum,* constituting different cluster sizes, shape and texture (Bamboo Technical Support Group (BTSG) 2025). However, we did not find any cavities in most of those bamboo species. Similar to us, *Tylonycteris* bats were found only in cavities of one bamboo species, *Gigantochloa scortechinii,* in Malaysia (Medway and Marshall 1970), and the authors report that there were no instances of beetle attack in any of the five other bamboo species in their study site. These differences suggest that cavity availability for bats is likely related to bamboo species and even cluster density, as insects might prefer certain species or cluster densities to complete their life cycle. Understanding the insect preferences and cavity formation dynamics requires systematic investigation, as it is relevant in the context of listing out potential bamboo species that are critical for bamboo bats and also to document the vulnerability of different bamboo species to pest infestation in this habitat.

### Cavity entrance and occupancy

Cavity mouth was the only factor associated with occupancy in our study, and cavity width was the only significant factor differing between the roosts of two sympatric *Tylonycteris* species in Malaysia (Medway and Marshall 1970). Under the sympatric condition, the larger *T. robustula* excluded the narrowest cavities, suggesting an ecological separation based on this one roost property. Thus, the selection of a cavity for roosting is likely dictated by the cavity and body size compatibility of bamboo bats. Cavities with a certain mouth area might be best suited for occupancy among all the available cavities in our case as well. However, we had measurements from only 21 unoccupied cavities, which is not a complete representation of the actual size range of all the available cavities in the habitat for bats to occupy at a given time. Having such a threshold size range is still plausible, as it can be advantageous for the bats in multiple aspects.

The cavity mouth is the point of entry and exit, and the smallest mouth area of an occupied cavity in our dataset was 0.87 cm^2^. It is possible that cavities with mouth areas below this can make it hard for the bats to enter and exit. However, bamboo cavities are utilised by a range of cavity-dwelling taxa in this landscape, and smaller cavities might still be ideal for smaller cavity-dwelling insects and arachnids (Kovac and Streit 1996). Similarly, larger mouth areas might aid easy entrance and exit for bats but can also provide accessibility for other cavity-dwelling taxa and predators. Thus, bats might need to defend their roosts from both competitors and predators or even abandon cavities above a certain mouth area. Our observations hint towards the possibility of such a cavity size threshold, as the roost with the largest mouth area (3.16 cm^2^) was also one of the three abandoned cavities within the course of our study.

Interestingly, the second abandoned cavity was found occupied by a spider (presumably a huntsman spider) through CCTV observations. We noted that the spider came out of the cavity, and a bat was flying at the cavity entrance while the spider was already present. The spider moved away from the cavity mouth after multiple such circling flights by the bat towards it (Supplementary video 1). However, the bat did not enter inside after that, and the cavity was confirmed to be subsequently abandoned. Another huntsman spider was found at a cavity entrance right next to a solitary roost from a different cluster. In addition, we observed centipedes, snakes, small invertebrates, and geckos active either near or at the cavity entrances (Supplementary Fig. 1). These observations were not restricted to any single bamboo cluster and thus support the possibility of bamboo bats preferring cavities with a particular mouth area beyond which it might become unsuitable for roosting. However, a robust sampling design that includes all available cavities and continuous monitoring of occupied and unoccupied cavities is required to investigate this thresholding better.

### Seasonality and roosting dynamics

One limitation of our study is that we collected information on occupancy in a limited time frame and assigned roost identity based on that. While this is relevant in terms of understanding how physical properties of the bamboo cavities are associated with roosting at a given time point, it does not provide insights on the factors influencing continued occupancy. Except for the mouth area, the properties of the unoccupied cavities did not differ much from those of occupied cavities, and even the mouth area of unoccupied cavities was within the range of occupied ones (Fig. 2b). This suggests that cavity mouth area is likely the parameter influencing occupancy at a given time point but might not be a stand-alone factor determining continued occupancy. With the current dataset, we do not know whether those unoccupied cavities were once occupied or can become occupied in the future. Similarly, the certainty of the current occupied cavities being continuously occupied is also unclear from the present design. It is also possible that bats switch cavities across seasons, as known in other *Tylonycteris* species (Medway and Marshall 1972) and might prefer larger-mouthed cavities when they are pregnant or lactating to aid easier entries and exits. Thus, occupancy in itself is possibly dynamic and likely varies across seasons. Roost preference thresholds, if any, might also change with temporal factors in addition to physical parameters. Continuous monitoring of cavities, along with the dynamics of cavity formation, growth, and occupancy status across time, is essential to understand this system in further detail.

### Roost properties and group sizes

By observing the roosting associations in two sympatric bamboo bat species, *T. pachypus* and *T. robustula,* in Malaysia, Medway and Marshal (1972) clearly pointed out that the number of bats in a roost is limited by factors other than roost characteristics. Our results also support this proposition (Fig. 4). Beyond available space, other factors such as sex ratio, sociality, mate choice, and even breeding status can shape the composition of bamboo bat groups. Group composition was similar in *T. pachypus* and *T. robustula* across the year in Malaysia, with solitary bats being most frequent, accounting for 36 and 52% of all roosting groups, respectively. Among these, solitary males were more common, but there were cases of solitary females as well (Medway and Marshall 1972). We found 26% of our occupied cavities were solitary, but we could not confirm whether they were males or females during the study period. Following solitary, both *T. pachypus* and *T. robusta* had group sizes ranging from pairs to up to 20 and 32 individuals, comprising both unisexual and mixed sex groups over the year. However, seasonal variations linked to reproduction have also been reported in both species. Males are reported to become reproductively active in October, followed by mating, and females are in oestrus until January. Our study was conducted in November-December and falls within this season, when probably both males and females were reproductively active. During November and December, *T. pachypus* males in Malaysia were found either to be solitary or associated with females but not in pairs or male-only groups (Medway and Marshall 1972). Interestingly, during a bat photography session conducted in January 2026 at one of our solitary roosts, we noticed a male bat flying out, confirmed by the presence of a pair of testicles visible through the camera (authors pers.obs). Being a closely related species, if not the same species, it is likely that all our solitary roosts also had only males in November, but this needs further investigation. Capture studies with individual marking using RFID tags, continuous monitoring of roosts, supplemented with endoscopic observations within the internodes whenever possible, can aid in recording sex ratios, group composition and seasonal variations in social organisations within these identified roosts.

### Ephemerality of bamboo roosts

With adaptations such as the flattened skull, thumb pads and soles in the feet, *Tylonycteris* species are specialised for roosting inside bamboo internodes, and this makes them completely dependent on one substrate for roosting without alternatives. However, bamboo is also one of the commonly harvested crops, as the culms serve as a commercial resource for furniture, handicraft and even for construction (Maurya et al. 2025; Anitha et al. 2026). Some species are also referred to as ‘green gold’ and ‘poor man’s timber’, as they are low-cost alternatives to timber (Chongtham et al. 2011). In addition, mature bamboo is also vulnerable to climatic changes, as it can easily uproot after heavy winds and rains. One of our three abandoned cavities was from one such bamboo culm that fell after winds. From the bats’ perspective, these aspects already place bamboo as a resource with high demand and low stability over time. The insects that make cavities in bamboo are also considered pests and sometimes controlled (Joshi et al. 2008). Loss of these insects would mean less availability of habitable roosts and, on the other hand, an increase in competition among other cavity-dwelling taxa. The increasing demand for bamboo, unsustainable harvesting practices (India State of Forest Report 2021; Bamboo Technical Support Group (BTSG–KFRI)), along with possible loss of the cavity-boring insects, not only threatens the bamboo bats in this region but also makes their roost markedly ephemeral. We emphasise the importance of these ephemeral bamboo cavities to the elusive and rare bamboo bats in this region and highlight the need to develop sustainable bamboo management strategies to conserve biodiversity and support ecosystem health.

As the bamboo cavities become more ephemeral, the bats are also in constant need of switching roosts and searching for more habitable cavities frequently. Unlike other bat species that have stable roosts, ephemeral roosting bats need to invest time in both roost finding and foraging during the nights. An extreme example of ephemeral roosting followed by roost switching is the Spix’s disk-winged bat *Thyroptera tricolor, which* roosts in the furled leaves of *Heliconia* and *Calathea* plants in the Neotropics (Findley and Wilson 1974; Vonhof and Fenton 2004). With suction disks in the wrists and ankles, *T. tricolor* are also specialised to roost in an unstable substrate and switch roosts daily (Riskin and Fenton 2001; Vonhof and Fenton 2004). Despite this, they are still known to be successful in finding new roosts, form stable long-term associations among individuals and have specialised behavioural strategies to regroup through complex social communication systems after being separated (Chaverri et al. 2010; Chaverri 2010; Montero and Gillam 2015). Similar behavioural strategies, social communication and long-term associations are highly likely in *Tylonycteris* bats, as they are also presented with similar challenges as *T. tricolor.* However, we lack any knowledge on any of these aspects in bamboo bats, and our results provide the preliminary ecological framework with respect to roost selection and group sizes that can assist in designing informed behavioural studies.

### Conclusion

There is still taxonomic ambiguity in the species found in this region, and recent multivariate analysis of this population suggests this to be a potential new species (Bates and Harrison 1997; Tu et al. 2017; Wilson and Mittermeier 2019; Raman and Hughes 2020; Raman et al. 2021; Chelladurai 2025). An integrative taxonomic approach is essential to resolve the species name to promote a site- and species-specific threat assessment and informed conservation strategies. Irrespective of this being a new species or a subspecies of a different population, this is the only bamboo bat species known in the Western Ghats. The overall habitat, threats, and even the specific insect and bamboo species associated with this bat species are likely unique from other *Tylonycteris* species or populations. We still lack knowledge of most of the critical ecological aspects of this rare and elusive bat species that are highly dependent on the tropical bamboo forests (Raman et al. 2021, 2022). Our findings provide first-hand knowledge on the roosting ecology of these bats and insights into this understudied ecological continuum that involves bats, bamboo, insects, cavity-dwelling taxa and bamboo harvesters in the Western Ghats of India. Overall, these findings have also opened up key future avenues with respect to bamboo bat ecology, behaviour, bamboo pest dynamics, threat assessment and finally conservation of the less known ecological continuum in the Indian tropics.

## Acknowledgements

We thank the Kerala Forest and Wildlife Department for granting permission for studying bats in various landscapes of Kerala. KSCSTE - Kerala Forest Research Institute provided organisational and infrastructural support for this study. We are grateful to Krishnapriya, Botany Department, KFRI, for inputs and help identifying the bamboo species reported in the study. SR was supported by the Merlin Tuttle’s Bat Conservation through the Antoniya Hubancheva Award during this study. We sincerely thank Daniel Hargreaves and Melissa Donnelly for their valuable inputs and for sharing the images used in this manuscript.

## Author contributions

SR: Conceptualisation, methodology, investigation, resources, writing – original draft, writing – review and editing, funding acquisition

BM: Conceptualisation, formal analysis, investigation, validation, visualisation, writing – original draft, writing – review and editing

MRJ: Methodology, data collection, data curation, formal analysis, writing – original draft, writing – review and editing

AJ: Methodology, data collection, data curation, writing – review and editing UVA: Methodology, data collection, data curation, writing – review and editing

PB: Project administration, Investigation, resources, supervision, writing – review and editing

## Data availability

All cavity morphometrics data used in this manuscript have been included in the supplementary section as Supplementary Table 2

## Supplementary information

*Supplementary file*: This file contains supplementary figures and tables supporting our study.

*Supplementary Table 2*: Excel sheet containing raw data of all roost morphometrics used in the study. A descriptive README file is included as sheet 1 for further clarity.

*Supplementary video 1*: Video showing a spider coming out of an occupied cavity. The spider remained at the cavity mouth, during which a bat flew around it until the spider moved away. This cavity was first occupied but subsequently confirmed as abandoned, as there were no bats inside after the spider was spotted.

## Compliance with ethical standards

### Permits and ethical guidelines

This study reports findings from our original research. No animals were captured, handled or harmed during this study. This is strictly an observational study following the permitted national guidelines. Permissions for conducting research on bats in this study landscape were obtained from the Kerala Forest Department (permit number_KFDHQ/4818/2025-CWW/WL-10 & KFDHQ-4009/2021-CWW/WL-10).

