## Supplementary file for "Hollow homes: Roost selection and role of roost morphometrics in the cavity-roosting Bamboo Bat (Genus *Tylonycteris*) from the Western Ghats, southern India"

### ELECTRONIC SUPPLEMENTARY FILE

**Supplementary Table 1:** GLM summary of the global model for roost preference. Colony size (solitary or non-solitary) was the binary response variable. Inclination angle, cavity mouth area, height from the ground, and roost volume were predictors.

| Predictor | Odds Ratio | CI Lower | CI Upper | Std. Error | z value | p value |
|---|---|---|---|---|---|---|
| Intercept | 0.083 | 0 | 5055.627 | 5.121 | -0.487 | 0.626 |
| Cavity mouth area (cm²) | 2.222 | 0.147 | 58.409 | 1.434 | 0.557 | 0.578 |
| Roost volume (L) | 72.938 | 0.322 | 261038.331 | 3.268 | 1.313 | 0.189 |
| Height from ground (m) | 1.181 | 0.289 | 4.789 | 0.665 | 0.25 | 0.803 |
| Inclination angle | 0.971 | 0.86 | 1.082 | 0.054 | -0.553 | 0.581 |

**Supplementary Table 2:** Mann-Whitney U statistics for the four tested predictors when compared between solitary and non-solitary roosts

| Predictor | W statistic | p value |
|---|---|---|
| Cavity mouth area (cm²) | 41 | 0.611 |
| Roost volume (L) | 19 | 0.151 |
| Height from ground (m) | 24 | 0.331 |
| Inclination angle (°) | 43.5 | 0.458 |

**Supplementary Figures**

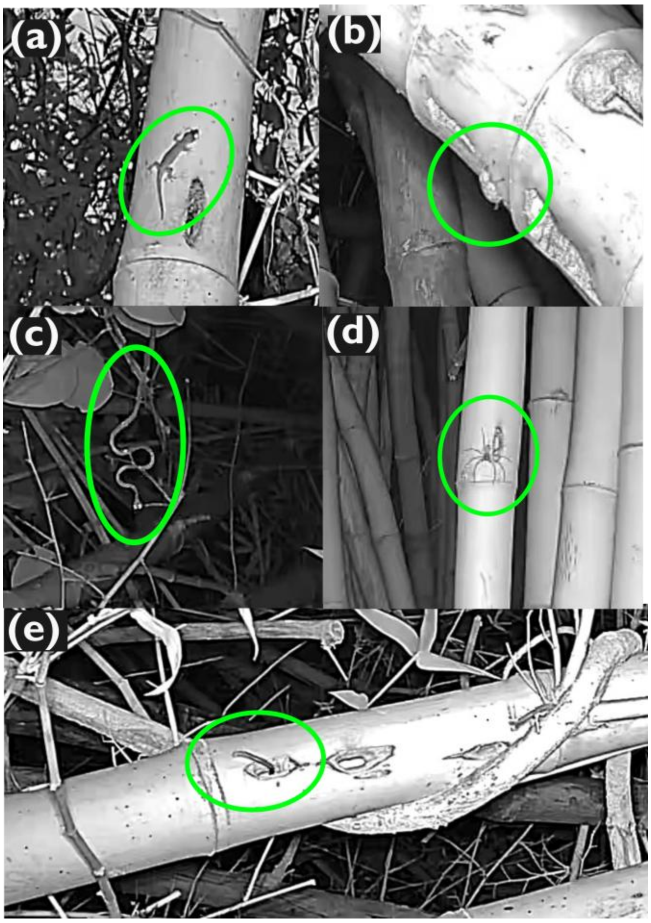


**Supplementary Figure 1: Cavity-dwelling taxa found active around roosts of *Tylonycteris pachypus aurex.*** (a) gecko, (b) beetle, (c) snake, (d) spider, (e) centipede. All these observations are based on CCTV footage from the roosts during our study period.
